# Validation of Fourier transform infrared microspectroscopy for the evaluation of enzymatic cross-linking of bone collagen

**DOI:** 10.1101/2023.02.27.530291

**Authors:** Aleksandra Mieczkowska, Guillaume Mabilleau

## Abstract

Enzymatic cross-linking of the bone collagen is important to resist to crack growth and to increased flexural strength. In the present study, we proposed a new method for assessment of enzymatic cross-link based on FTIR microspectroscopy that takes into account secondary structure of type I collagen. Briefly, femurs were collected from sham or ovariectomized mice and subjected either to LC-MS or embedded in polymethylmethacrylate, cut and analyzed by FTIR microspectroscopy. FTIR acquisition were recorded before and after UV exposure or acid treatment. In addition, femurs from a second animal study were used to compare gene expression of *Plod2* and *Lox* enzymes and enzymatic cross-links determined by FTIR microspectroscopy.

We evidenced here that intensities and areas of subbands located at ∼1660 cm^-1^, ∼1680 cm^-1^ and ∼1690 cm^-1^ were positively and significantly associated with the concentration of pyridinoline (PYD), deoxypyridinoline (DPD) or immature dihydroxylysinonorleucine (DHLNL) / hydroxylysinonorleucine (HLNL) cross-links. Seventy-two hours exposure to UV light significantly reduced by ∼86% and ∼89% the intensity and area of the ∼1660 cm^-1^ subband. Similarly, 24 hours of acid treatment significantly reduced by 78% and 76% the intensity and area of the ∼1690 cm^-1^ subband. *Plod2* and *Lox* expression were also positively associated to the signal of the ∼1660 cm^-1^ and ∼1690 cm^-1^ subbands.

In conclusion, our study provided a new method for decomposing the amide I envelope of bone section that positively correlates with PYD and immature collagen cross-links. This method allows for investigation of tissue distribution of enzymatic cross-links in bone section.

## 1. INTRODUCTION

The extracellular matrix of bone tissue is a composite material, consisting of type I collagen fibres (organic matrix, ∼35-45 % by volume), poorly crystalline carbonated hydroxyapatite crystals (mineral phase, ∼35-45% by volume) and water (∼15-25% by volume) ^(1)^. Although type I collagen is the most ubiquitous protein in connective tissues, its chemistry varies from one tissue to another due to post-translational modifications and cross-linking ^(2)^. Collagen cross-linking add stability to the organic matrix, preventing the microfibrils from sliding over each other, allowing the cortical bone to better resist to crack growth and to increase bending strength of long bones ^(3)^.

Enzymatic collagen cross-linking occurs in the extracellular space, however, the nature of the cross-links depends on previous intracellular post-translational modifications of the collagen molecule ^(4)^. The first step of enzymatic cross-linking is represented by the intracellular hydroxylation of lysine (Lys) residue within the triple helix by lysyl hydroxylases 1 and 3 ^(5)^ and within telopeptides by lysyl hydroxylase 2 ^(6)^ leading to the formation of hydroxylysine residue (Hyl). In the extracellular space, Lys and Hyl residues can be converted to aldehydes, respectively Lys^ald^ and Hyl^ald^, by the action of lysyl oxidase. These aldehydes then initiate a series of condensation reactions with vicinal Lys^ald^, Lys, Hyl and histidine residues to form intra and intermolecular covalent cross-links ^(7)^. In bone, the predominant enzymatic cross-links found are the divalent immature cross-links, dehydrodihydroxylysinonorleucine (deH-DHLNL) and dehydrohydroxylysinonorleucine (deH-HLNL) ^(8)^. These immature cross-links are unstable, destroyed by acid treatment ^(9)^ and need to be reduced into their stable counterparts, dihydroxylysinonorleucine (DHLNL) and hydroxylysinonorleucine (HLNL), in order to be detected by LC-MS. Mature trivalent cross-links predominant in bone tissue are hydroxylysylpyridinoline (PYD), lysylpyridinoline (DPD) and pyrroles (Pyr).

The gold standard for the analysis of collagen cross-links is represented by LC-MS ^(10)^. However, LC-MS techniques are destructive and require bone samples to be powdered prior to analysis. With such setup it is not possible to evaluate the distribution of collagen cross-links within a tissue despite recent evidences suggest that heterogeneity in collagen cross-link content is key for higher bone resistance to fracture ^(11)^. Twenty-two years ago, Paschalis et al. proposed a spectroscopic method for the evaluation of collagen cross-links in bone tissue sections ^(12)^. This methodology is based on Fourier transform infrared (FTIR) microspectroscopy analysis, in transmission mode, of the amide I and II spectral regions after second derivative and curve fitting. Paschalis et al. evidenced that two subbands located at ∼1660 cm^-1^ and ∼1690 cm^-1^ corresponded to perturbation of carbonyl group vibrations due to PYD and deH-DHLNL ^(12)^. More recently, Paschalis et al. also showed that not only the ∼1660 cm-1 subbands was indicative of trivalent collagen cross-links but also the ∼1680 cm-1 subband that seemed correlated to perturbation of carbonyl group vibrations due to the presence of DPD ^(13)^. The 1660/1690 cm^-1^ ratio is now widely used worldwide as a marker of collagen maturity and represents an index of the maturation of divalent to trivalent cross-links. The 1660/1680 cm^-1^ is also more and more used as an indicator of enzymatic collagen cross-link ratio ^(14)^. However, differences in FTIR spectral processing as compared with the original method reported by Paschalis et al., are often found and the validity of such maturity and cross-link ratios remains to be fully validated against gold-standard LC-MS method. Recently, the ∼1678/1690 cm^-1^ ratio has also been proposed for investigation of advanced glycation end products in bone ^(15)^. As such, better assignment of the subband located ∼1680 cm-1 is required to appreciate the validity of newer outcome.

A second point of attention resides in the method used to identify subbands in Amide I spectral region. Indeed, by rigidly imposing the presence of subbands at 1633, 1660 and 1690 cm^-1^, Farlay et al. were incapable of recording any significant decrease in enzymatic collagen cross-linking between controls and β-aminopropionitrile-treated animals ^(16)^. However, when enzymatic collagen cross-links were measured by LC-MS, a significant reduction in PYD and DPD was observed between the two groups of animals. Interestingly, Belbachir et al. reported previously the presence of 9 subbands in the amide I band related to the secondary structure of type I collagen ^(17)^. Moreover, 4 additional subbands could be present due to amino acid side chains ^(18)^. In its seminal paper investigating enzymatic collagen cross-links by FTIR spectroscopy, Paschalis et al. described only 4 subbands in the amide I envelope of demineralized bovine bone ^(12)^. More recently, Schmidt and coworkers reported their method of amide I decomposition that used seven gaussian subbands located between 1600 cm^-1^ and 1710 cm^-1 (15)^. The spectral signatures of enzymatic collagen cross-linking overlap in the amide I band with spectral signature of the secondary structure of type I collagen and amino acid side chains. As such, it seems important to develop and validate a new method for decomposing the amide I signal that take into account the secondary structure of type I collagen and amino acid composition in order to accurately measure reducible divalent and non-reducible trivalent collagen cross-links. The accuracy of divalent and trivalent subbands needs also to be validated against gold standard LC-MS methodology for biological assignment.

In the present study, we reported a new second derivative spectroscopy-curve fitting algorithm based only on amide I spectral region and taking into account secondary structure of type I collagen and amino acid side chain for the determination of enzymatic collagen cross-links in bone tissue. Our method was compared with LC-MS determination of collagen cross-link content and the extent of gene expression of lysyl hydroxylase 2 and lysyl oxidase, two key enzymes involved in the initiation of enzymatic cross-linking.

## 2. MATERIAL AND METHODS

### 2.1. Animal models

All procedures were carried out in accordance with the European Union Directive 2010/63/EU for animal experiments and were approved by the regional ethical committee for animal use (authorization CEEA-PdL06-01740.01). Evaluation of enzymatic collagen crosslinking was realized in bones of two animal studies. Figure 1 represents a schematic of the experimental protocol. In the first study, 20 BALB/c female mice (BALB/cJRj) were purchased from Janvier Labs (Janvier Labs, St Berthevin, France). Ten animals underwent Sham surgery at 12 weeks of age and the remaining ten animals underwent bilateral ovariectomy under general anesthesia at the same age. Animals were sacrificed 4 weeks post-surgery. These animals were used for the correlation between collagen crosslinks concentrations by HPLC-MS and FTIR microspectroscopy. In the second study, bilateral OVX was performed in 8 BALB/c (BALB/cJRj) female mice at 12 weeks of age under general anesthesia. Eight sham operated BALB/c female mice with the same age were also used. These animals were used for the correlation between gene expression of *Plod2* and *Lox*, 2 key enzymes involved in enzymatic collagen crosslinking ^(19,20)^. All animals were housed in social groups and maintained in a 12 h:12 h light:dark cycle and had free access to water and diet. At necropsy, uterus were collected and weighted to ensure optimum ovariectomy. Both femurs were also harvested and cleaned of soft tissues. Epiphyses were removed and the bone marrow was flushed out with PBS (Study 1) or removed by centrifugation (Study 2) as proposed by Kelly et al. ^(21)^. After bone marrow removal, left femurs were snap frozen in isotonic saline (Study 1) or RNAlater (Study 2) and store at -80°C until use.

**Figure 1:**
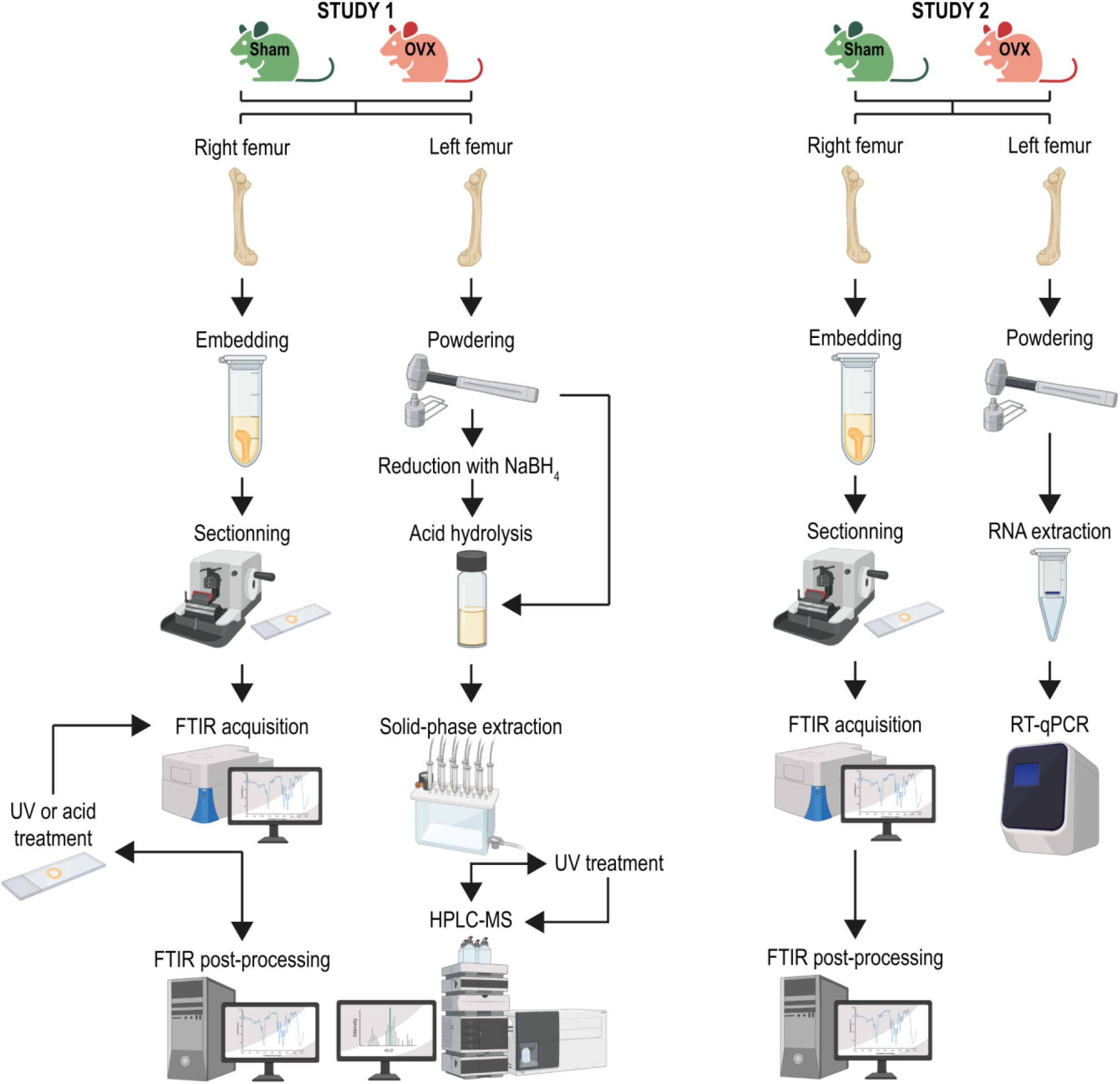
Schematic of the study. In study 1, right and left femurs from 10 Sham and 10 OVX mice were collected, cleaned of soft tissue and process for either FTIR or HPLC-MS analyses. In study 2, right and left femurs from 8 Sham and 8 OVX mice were collected, cleaned of soft tissue and process for either FTIR or gene expression analyses.

### 2.2. Processing of bone samples for the evaluation of collagen crosslinks by FTIR analysis

After overnight thawing at 4°C, the right femur was fixed with 70% ethanol, dehydrated, cut transversally in half at the midpoint between third trochanter and distal condyle with a diamond saw, and embedded undecalcified in pMMA. One micrometer-thick cross-section of proximal and distal part of the same femur was cut with an ultramicrotome (Leica EM UC7, Leica microsystems, Nanterre, France) and deposited on BaF2 windows. Spectral analysis was performed with a Bruker Hyperion 3000 infrared microscope coupled to a Vertex 70 spectrometer using a 64 × 64 focal plane array detector. A field of view of 540 × 540 µm covering the posterior quadrant of the femur midshaft was analyzed. The spectrometer was filled with dessicant pellets to reduce the contribution athmospheric water and CO_2_. A 15X Cassegrain objective (NA 0.4) was used for all acquisition. The FPA detector was cooled down with liquid nitrogen for higher sensitivity. Mid-infrared spectra were recorded at a resolution of 4 cm^-1^ (spectral region 900-2000 cm^-1)^, with 32 accumulations in transmission mode. Background spectra were also recorded with the same specifications. Post-processing of spectra was done with a lab-made script written in Matlab and includes Mie scattering correction ^(22)^, pMMA subtraction, normalization of v1,v3 PO4 peak to 1 and denoising using the Savitzky-Golay algorithm with a degree of 2 and a span length of 9. A quality control of each spectrum was performed by calculating the signal-to-noise ratio (SNR) over the spectral range 1850-2000 cm^-1^ free of biological signal. The SNR was computed as the ratio between the mean intensity on the spectral range 1850-2000 cm^-1^ over the standard deviation in the same spectral range. A SNR ≥ 10 was judged sufficient to continue the post-processing. Bone pixels were manually thresholded from bone marrow and resin using the v1,v3 PO_4_ peak area. Non bone pixels were assigned as NaN. Spectra were further subjected to second derivative of the Amide I band (1590 – 1730 cm^-1)^ for identification of subband locations. With this method, the band constituting a given interval are revealed by the successive minima of the second derivative curve. For simplicity and automation in detection of subbands, we inverted the second derivative to identify successive maxima. Band assignment is presented in Table 1. Validation of subbands was based on secondary structure of type I collagen and signal occurring from perturbation of carbonyl group vibration at ∼1660 and ∼1690 cm^-1^. Subband locations and widths at half maximum were then entered into a lab-written script that use a gaussian function for subband shape and allow the positioning with ± 3 cm^-1^ as compared with the location evidenced on the second derivative. Curve fitting was then processed to study separately the absorption bands of samples that may overlap in the amide I enveloppe. Spectral curve fitting quality was assessed by a root mean square error value set at 1% of the total spectral interval area. Intensities and areas of each subband were obtained. An example of the consequences of post-processing, second derivative and curve fitting on FTIR spectra is presented in Figure 2. Interassay variability, measured on five consecutive acquisitions of hyperspectral cubes on the same field of view in five consecutive days, implying repetition of internal calibration of the FTIR spectrometer and detector cooling as well as post-processing of the hyperspectral cubes, was of 0.2%.

**Table 1:**
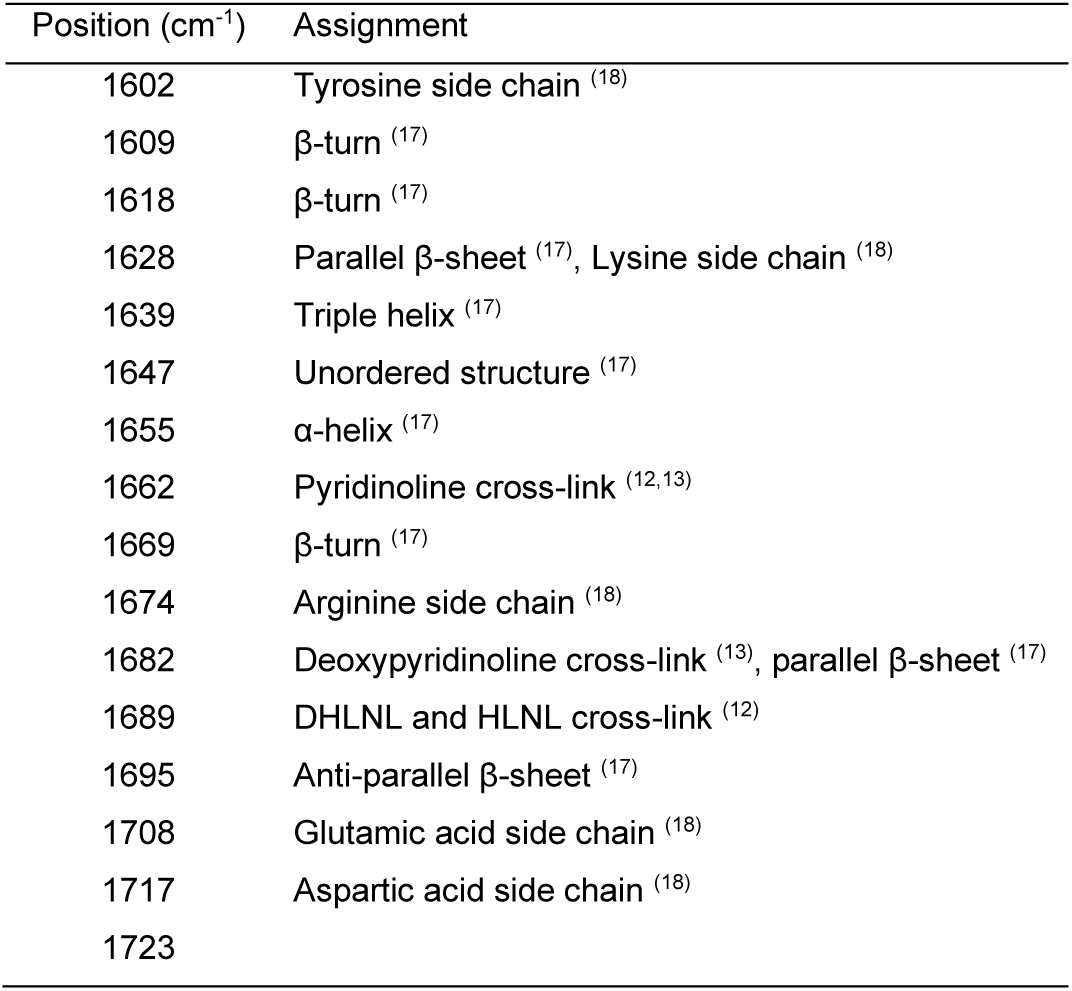
Subpeak position and assignment

**Figure 2:**
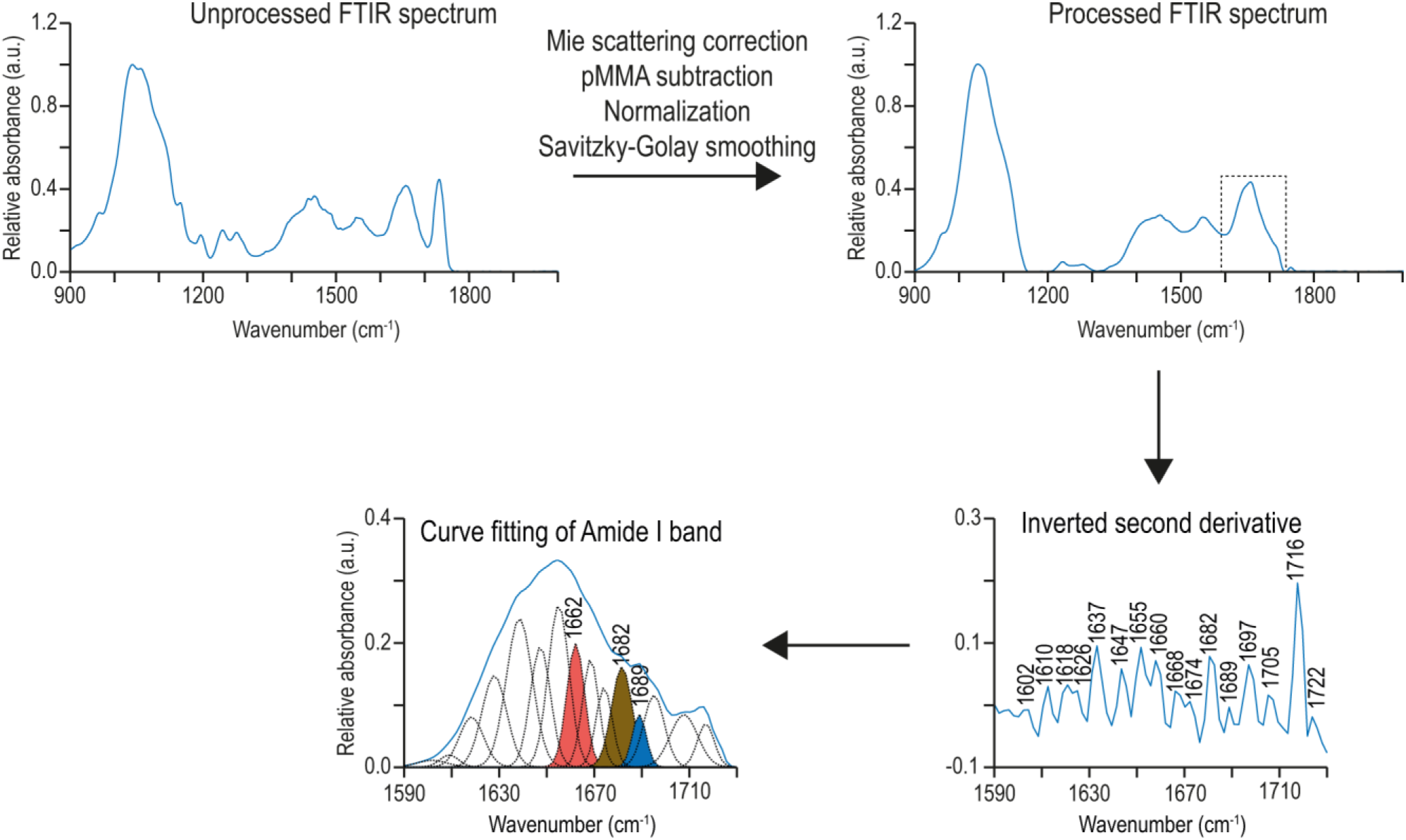
Example of FTIR post-processing. FTIR sample acquired directly from the bone section presented peaks due to biological tissue but also peaks due to the pMMA embedding resin. Furthermore, the raw spectrum may contain some noise that would interfere with subpeak detection. In order to extract useful information, FTIR spectra were processed to remove Mie scattering and the contribution of pMMA resin prior to normalization and Savitzky-Golay smoothing. The signal-to-noise ratio (SNR) was computed on processed spectrum. When SNR was ≥10, the spectrum was subjected to second derivative computation. Sixteen underlying subbands were detected and their positions and widths were used as a loading vector for curve fitting. Three subbands were of particular interest and highlighted in pink (∼1660 cm^-1)^, light brown (∼1680 cm^-1)^ and blue (∼1690 cm^-1)^.

After first acquisition, bone sections of the proximal part of the femur were exposed to a UV light for 72h at 4°C using a mercury lamp and an Olympus U-MNU2 filter with an excitation at 360 nm. This treatment is known to degrade mature collagen crosslinks. Similarly, bone sections from the distal part were exposed to 0.05M acetic acid at 4°C for 24h. This treatment is known to degrade labile immature collagen crosslinks. Then proximal and distal section were placed back to the FTIR microscope and the same field of view as analyzed previously was subjected to second FTIR acquisition as described above.

### 2.3. Processing of bone samples for the evaluation of collagen crosslinks by HPLC-MS

Standards for pyridinoline (PYD, also known as hydroxylysylpyridinoline) and deoxypyridinoline (DPD, also known as lysylpyridinoline) were purchased from BOC Sciences, Inc. (catalog # 63800-01-1 and B2694-136861, respectively). Dihydroxylysinonorleucine (DHLNL) was purchased from Santa Cruz Biotechnology (Catalog # sc-207059B). Hydroxylysinonorleucine (HLNL) was purified from bovine tibia collected in a local slaughterhouse as described below for mouse bone. The left femur was frozen with liquid nitrogen and powdered using a multisample biopulverizer (Biospec, Cat. No. 59012MS), defatted in methanol/chloroform, extensively washed with deionized water and freeze-dried. The bone powder was then demineralized with 0.5 M EDTA pH 7.4 for 96h at 4°C, with renewal of the demineralization solution every day. The bone powder was then resuspended in phosphate buffer saline and half of the resuspended powder was reacted with sodium borohydride (10 mg/ml in 1 mM NaOH with a ratio reagent/bone powder of 1:30 (w/w)) for 2h at room temperature to preserve the immature collagen crosslinks ^(23)^. Reduced and non-reduced samples were then washed, freeze-dried and hydrolyzed in 6M hydrochloric acid at 110°C for 24h as proposed in Gineyts et al. ^(24)^. A portion of the acidic hydrolysate was used for hydroxyproline assay (HPLC assay Biorad). Values of collagen crosslinks were then normalized to collagen content assuming 14% hydroxyproline by mass in collagen. The remainder of the acidic hydrolysate was cleaned on an SPE column (SPE Chromabond crosslinks, Macherey-Nagel). Briefly, 400 µl of sample hydrolysate was added to 2.4 ml of acetonitrile in a glass vial. This solution was then transferred to a SPE column previously equilibrated with a wash buffer made of 8 volume of acetonitrile, one volume of acid acetic and one volume of deionized water. The SPE column was extensively washed four times with 2.5 ml of wash buffer and rinsed with 100 µl of deionized water. Collagen crosslinks were eluted with 600 µl of 1% heptafluorobutyric acid (HFBA) in HPLC glass vials. Sixty microliters of this eluate were then analyzed by LC-MS using an Alliance 2795 module (Waters, Guyancourt, France) equipped with a Waters 2487 UV detector and coupled to a Bruker ESQUIRE 3000+ ESI ion trap mass spectrometer equipped with an electrospray source (Bruker, Wissembourg, France) assisted by the HyStar software (Bruker Daltonics). Briefly, collagen crosslinks separation was performed using an Atlantis T3 reverse-phase column (3 µm, 4.6 × 100 mm) at 25°C as proposed by Gineyts et al. ^(24)^. The column flow rate was 1ml/min. After the analytical column, the flow was split in half between fluorescence (0.6 ml/min) and MS (0.4 ml/min). Solvent A consisted in 0.12% of HBFA in 18 ohms pure water (as proposed and optimized by Ginets et al.^(24)^) and solvent B was 50% acetonitrile. The column was equilibrated with 10% of solvent B prior to use and chromatographic separation was achieved with a gradient elution from 10% to 20% solvent B in 40 min. PYD and DPD were monitored for fluorescence emission at 395 nm and excitation at 297 nm. The mass analyses were performed in positive ion mode. The target ions were [M+H]^+^ at m/z 308 for DHLNL, 292 m/z for HLNL, m/z 429 for PYD and m/z 413 for DPD. The conditions were as follows: spray voltage of 4.5 kV, collision gas, He; collision energy amplitude, 1V, nebulizer and drying gas, N2, 7L/min; pressure of nebulizer gas, 30 psi; dry temperature, 340°C, m/z 100-1000. Sixty microliters of the collagen crosslinks eluate were also exposed for 72h to UV light (excitation at 360 nm) prior to analysis by LC-MS as reported above.

### 2.4. Gene expression analysis

Gene expression analysis was performed after crushing left femur in a liquid nitrogen-cooled biopulverizer. Nucleozol (Macherey-Nagel, Hoerdt, France) was added on top of the bone powder and total RNA were purified with Nucleospin RNA set nucleozol column (Macherey-Nagel) according to the manufacturer recommendations. Total RNA was reversed transcribed using Maxima first strand cDNA synthesis kit (Thermofisher Scientific, Illkirch-Graffenstaden, France). Real-time qPCR was performed using TaqMan™ Fast advanced master mix and TaqMan Gene Expression Assays for *Lox* (Mm00495386_m1) and *Plod2* (Mm00478767_m1). The *B2m* endogenous control (Mm00437762_m1) was used for normalization using the 2^-ΔCT^ method.

### 2.5. Statistical analyses

Linear regression between concentration of PYD, DPD, DHLNL and HLNL determined biochemically, *Plod2* or *Lox* expression and the intensity or relative area of the subpeaks located at ∼1660 cm^-1^, ∼1680 cm^-1^ and ∼1690 cm^-1^ were performed using GraphPad Prism 8. Pearson correlation coefficient was computed to assess the direction and magnitude of covariation. Modification of intensity or relative area of subpeaks located at ∼1660 cm^-1^, ∼1680 cm^-1^ and ∼1690 cm^-1^ before and after UV exposure or acidic treatment were compared using a paired t-test. Differences were considered significant when p< 0.05.

## 3. RESULTS

### 3.1. Positive correlation between biochemical and spectral measurements of enzymatic collagen cross-links

As represented in Figure 3 and as expected from previous work^(13)^, pyridinoline (PYD) content, determined by LC-MS, was positively correlated with the intensity (R=0.746, p<0.001) and area (R=0.743, p<0.001) of the ∼1660 cm^-1^ subband. Deoxypyridinoline (DPD) content was positively correlated with both the intensity (R=0.747, p<0.001) and area (R=0.847, p<0.001) of the ∼1680 cm^-1^ subband. Furthermore, dihydroxylysinonorleucine (DHLNL) content and hydroxylysinonorleucine (HLNL) content, were also positively associated with both intensity (R=0.638, p=0.003 and R=0.579, p=0.008, respectively) and area (R=0.631, p=0.003 and R=0.581, p=0.007, respectively) of the ∼1690 cm^-1^ subband.

**Figure 3:**
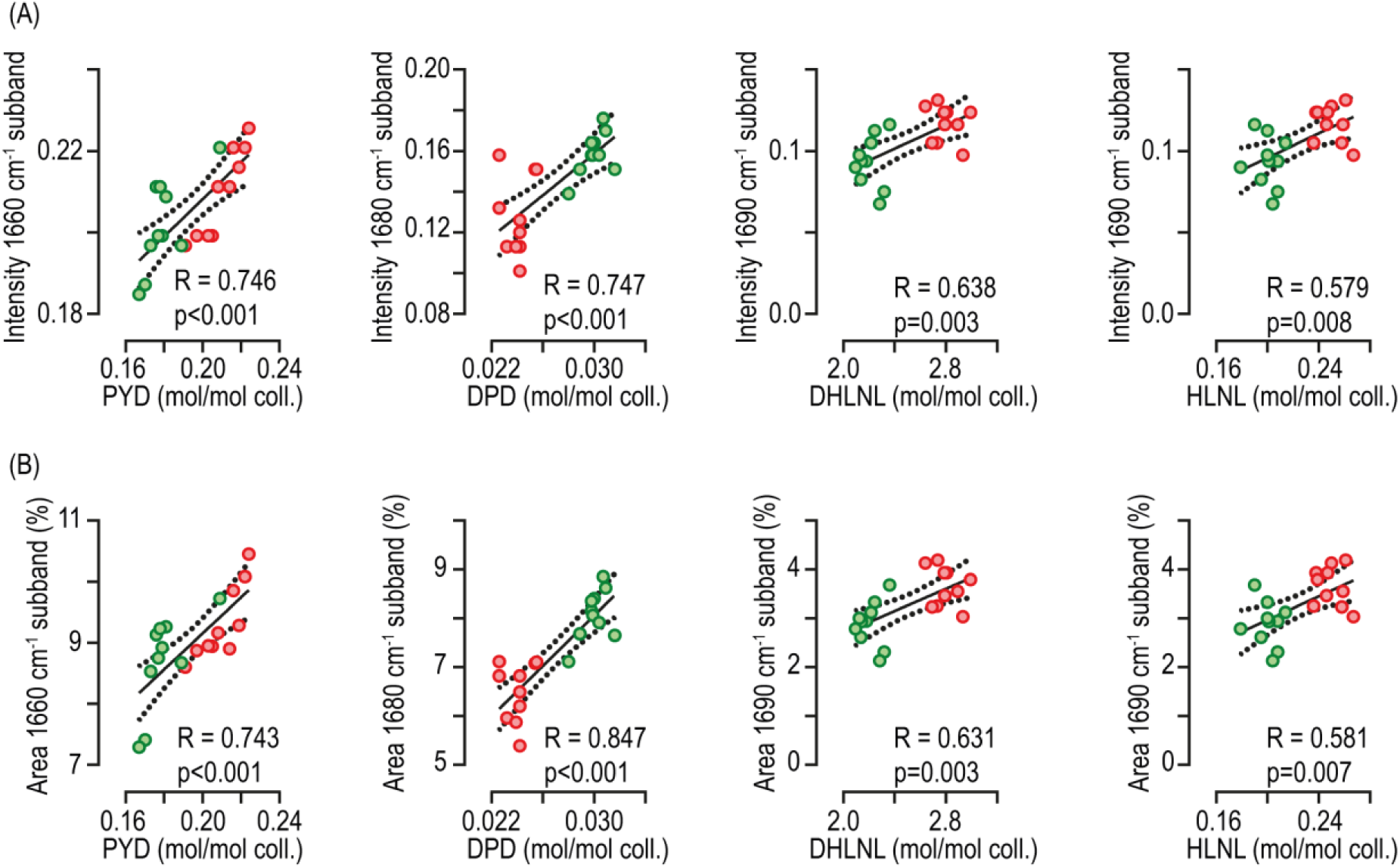
Correlations between concentration of collagen cross-links determined by LC-MS and intensity or area of the ∼1660 cm^-1^, ∼1680 cm^-1^ and ∼1690 cm^-1^ subbands. (A) Correlations between intensities of subbands and concentrations of pyridinoline (PYD), deoxypyridinoline (DPD), dihydroxylysinonorleucine (DHLNL) and hydroxylysinonorleucine (HLNL). (B) Correlations between area of subbands and concentrations of PYD, DPD, DHLNL and HLNL. The Pearson correlation coefficient (R) was computed and a two-tailed t-test was performed to assess the likelihood of random association between spectral and biochemical data, and considered significant at p<0.05.

### 3.2. Effects of UV exposure on mature collagen cross-links

Due to small sample size, the strong correlation presented above may also be purely due to chance. We then decided to study whether UV exposure, known to destroy mature collagen cross-links, may modulate intensities and areas of ∼1660 cm^-1^ and ∼1680 cm^-1^. Bone sections were acquired with the FTIR microscope before UV exposure. The same bone sections were then subjected to 72 hours of UV light and reacquired, at the same location, with the FTIR microscope using the same settings. A fraction of cross-link eluate was also subjected to UV exposure prior to LC-MS in order to validate that this treatment destroyed PYD and DPD. Indeed, PYD and DPD content, measured by LC-MS, were reduced by 91% (p<0.001) and 89% (p<0.001), respectively by this treatment (Figure 4A). Changes in amide I envelope was also observed after UV exposure (Figure 4B). Interestingly, intensity and area of the ∼1660 cm^-1^ subband were drastically reduced by 86% (p<0.001) and 89% (p<0.001) respectively, suggesting that the biological entity contributing to this spectral signal was altered by UV exposure (Figure 4C). On the other hand, intensity and area of the ∼1680 cm^-1^ subband were augmented by 46% (p<0.001) and 37% (p<0.001) respectively after UV exposure (Figure 4C). These results were in opposition to the reduction of DPD content evidenced by LC-MS.

**Figure 4:**
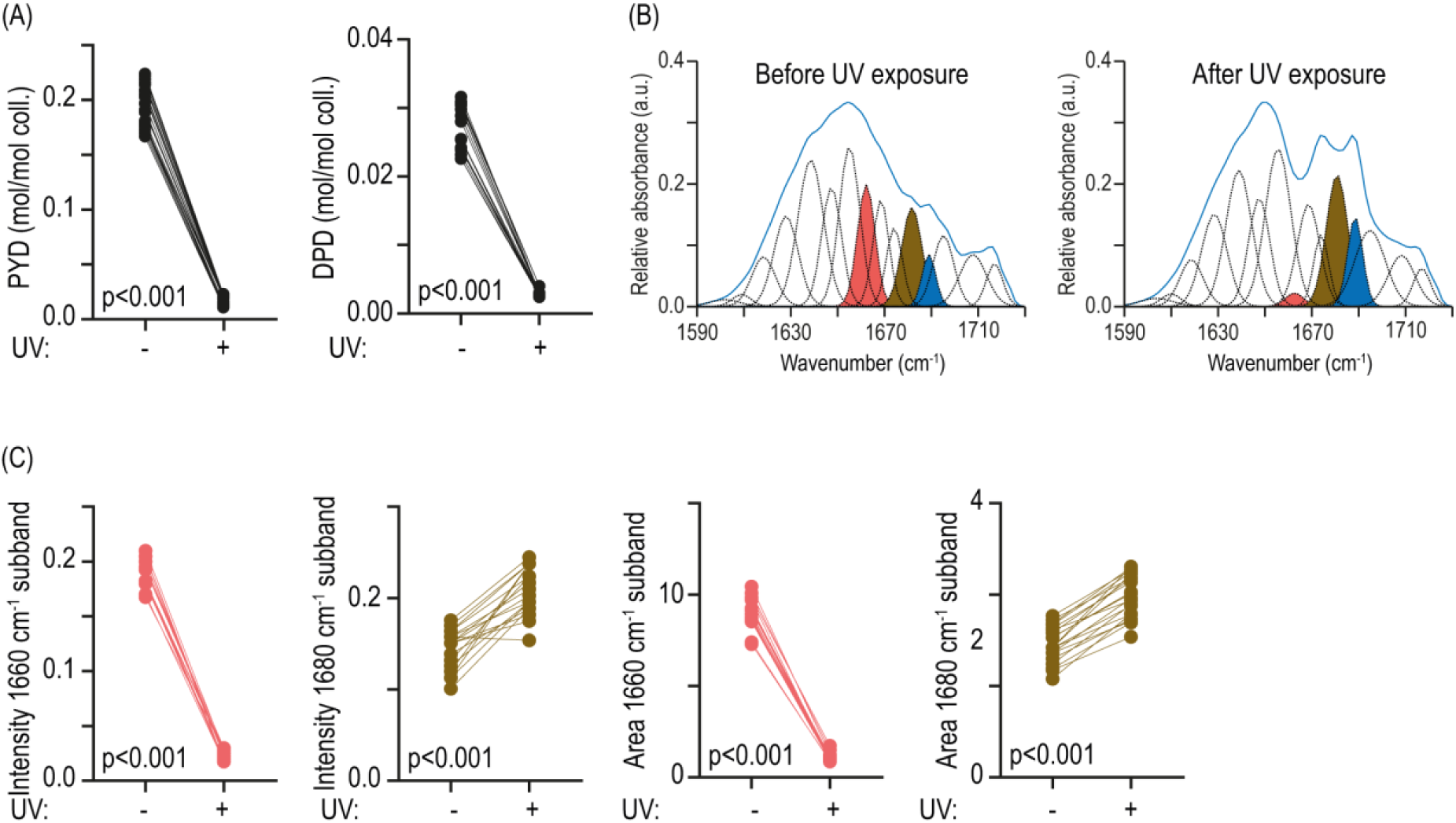
Effects of UV exposure of mature collagen cross-links. Mature collagen cross-links are sensitive to UV exposure. (A) PYD and DPD concentration were determined by LC-MS in two separate eluates that were exposed or not to UV light (360 nm) for 72 hours. (B) The same bone section was imaged by FTIR before and after UV exposure for 72 hours. Curve fitting of Amide I is presented and for clarity subbands located at ∼1660 cm^-1^, ∼1680 cm^-1^ and ∼1690 cm^-1^ were highlighted in pink, light brown and blue, respectively. (C) Intensities and areas of ∼1660 cm^-1^ and ∼1680 cm^-1^ subbands were computed on the same section prior to and after UV exposure for 72 hours. Two-tailed paired t-test were used to compute difference and were considered significant at p<0.05.

### 3.3. Effects of acid treatment on divalent collagen cross-links

Prior to acid hydrolysis, half of the bone powder was reduced by NaBH_4_ in order to reduce and protect divalent collagen cross-links. The remaining half was directly subjected to acid hydrolysis without reduction step in order to degrade divalent collagen cross-links. As represented in Figure 5A, the concentration of acid-labile divalent collagen cross-links was dramatically reduced by 99% (p<0.001) for DHLNL (or its native non-reduced deH-DHLNL) and by 99% % (p<0.001) for HLNL (or its native non-reduced deH-HLNL), respectively. Here again, treatment of the bone section by 0.05 M acetic acid for 24 hours resulted in changes in the amide I envelope (Figure 5B). Intensity and area of the ∼1690 cm^-1^ subband were significantly decreased by 78% (p<0.001) and 76% (p<0.001), respectively.

**Figure 5:**
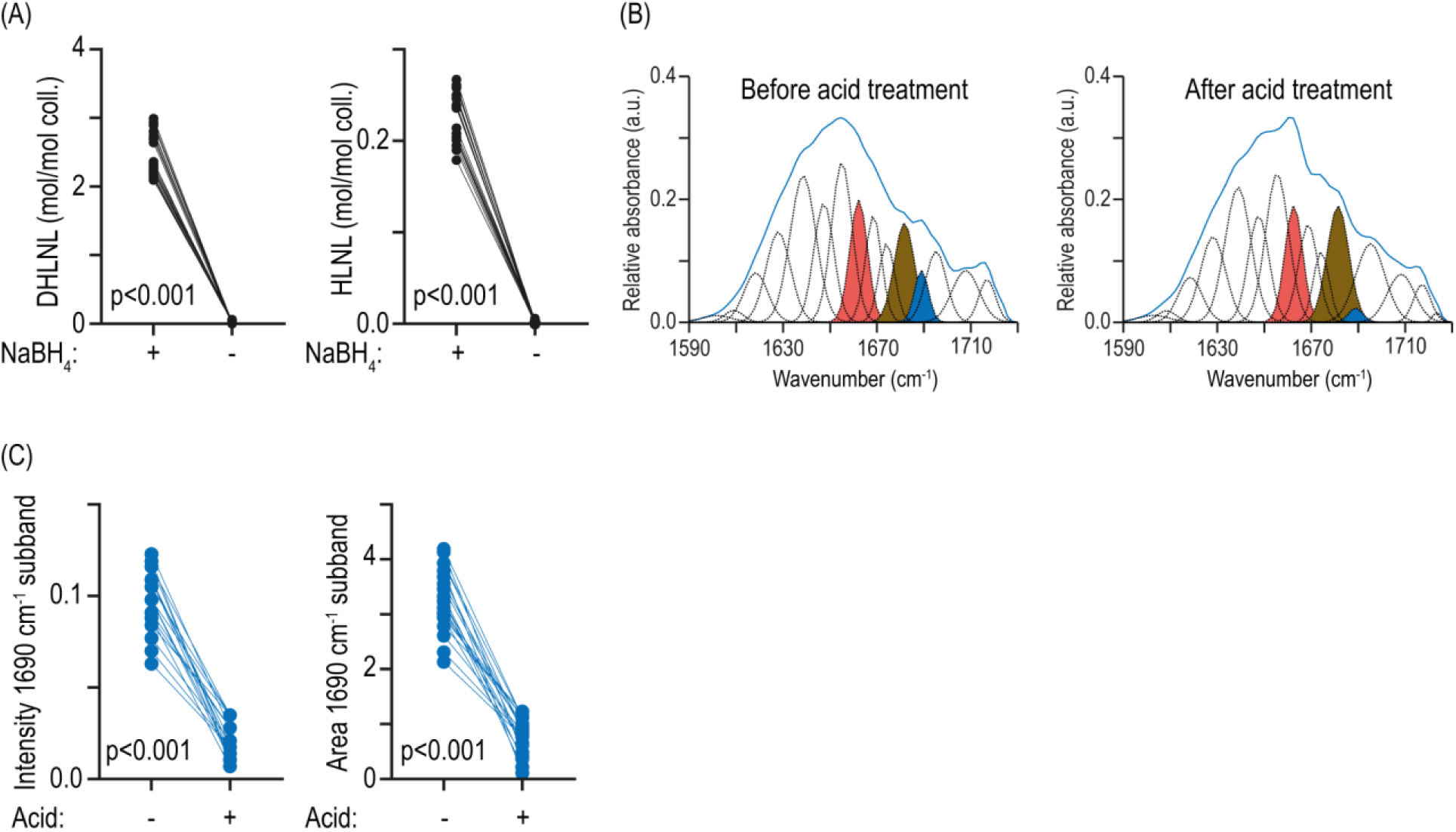
Effects of acid treatment on immature collagen cross-links. Immature collagen cross-links are unstable and destroyed by acid. (A) For LC-MS analysis, half of the bone powder was not reduced by NaBH4 prior to matrix hydrolysis in 6M HCl. The other half was reduced by NaBH_4_ and protected immature collagen cross-links for destruction. (B) The same bone section was imaged by FTIR before and after treatment with acetic acid 0.05 M for 24 hours. Curve fitting of Amide I is presented and for clarity subbands located at ∼1660 cm^-1^, ∼1680 cm^-1^ and ∼1690 cm^-1^ were highlighted in pink, light brown and blue, respectively. (C) Intensities and areas of ∼1690 cm^-1^ subband were computed. Two-tailed paired t-test were used to compute difference and were considered significant at p<0.05.

### 3.4. Correlation between expression of *Plod2* and *Lox*, and spectral measurements of enzymatic collagen cross-links

We next assessed in bone of another study whether gene expression of *Plod2* and *Lox*, two enzymes involved in lysine hydroxylation and oxidation required for initiation of divalent collagen cross-links, were correlated to the area of ∼1660 cm^-1^, ∼1680 cm^-1^ and ∼1690 cm^-1^ subbands. As presented in Figure 6, *Plod2* expression was significantly and positively correlated with area of ∼1690 cm^-1^ subband (R=0.548, p=0.028) and almost reach statistical for area of the ∼1660 cm^-1^ (R=0.487, p=0.056). However, no correlation was found for area of the ∼1680 cm^-1^ and *Plod2* expression (R=0.242, p=0.367). *Lox* expression was significantly and positively correlated with areas of ∼1660 cm^-1^ (R=0.502, p=0.048) and ∼1690 cm^-1^ (R=0.681, p=0.004) subbands. Here again, no correlation between *Lox* expression and area of the ∼1680 cm^-1^ subband could be observed (R=0.238, p=0.374).

**Figure 6:**
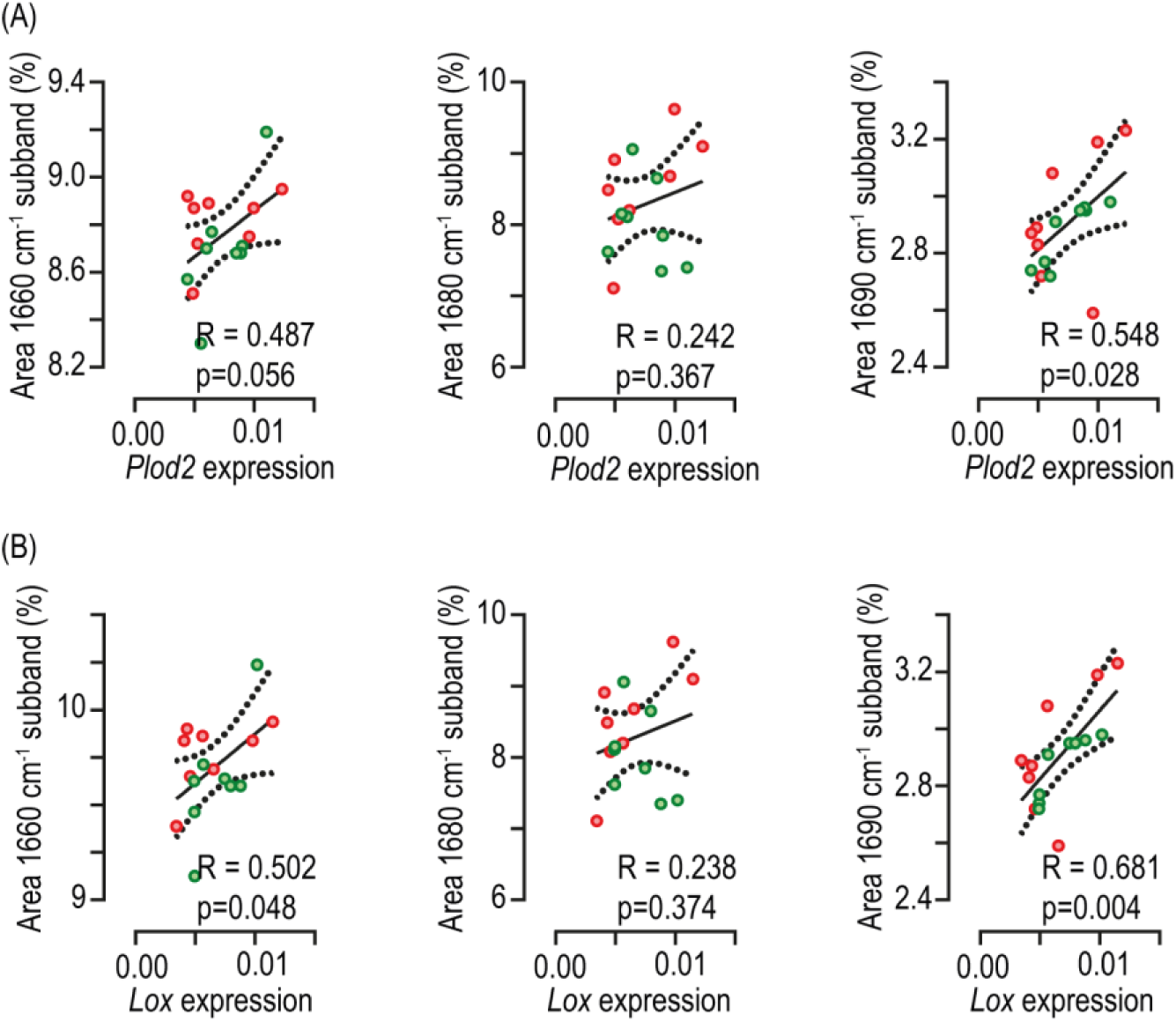
Correlations between gene expression of key enzymes involved in collagen cross-linking and area of the ∼1660 cm^-1^ and ∼1690 cm^-1^ subbands. (A) Correlations between level of *Plod2* expression and area of ∼1660 cm^-1^ and ∼1690 cm^-1^ subbands. (B) Correlations between level of *Lox* expression and area of ∼1660 cm^-1^ and ∼1690 cm^-1^ subbands. The Pearson correlation coefficient (R) was computed and a two-tailed t-test was performed to assess the likelihood of random association between spectral and biochemical data, and considered significant at p<0.05.

**Figure 7:**
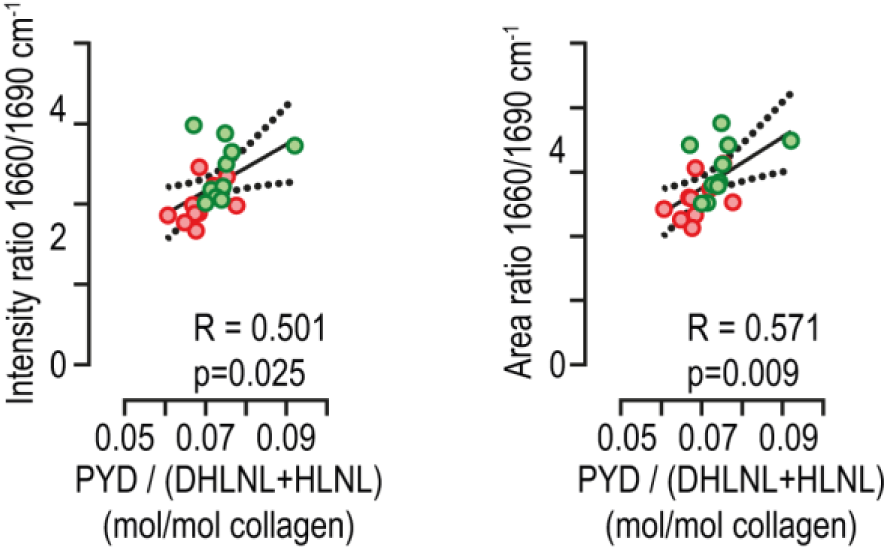
Correlations between biochemical and spectral collagen maturity ratios. The biochemical ratio was computed as the concentration of the mature PYD over the sum of DHLNL and HLNL concentrations. The spectral ratio was computed as the intensity or area ratio of subbands located at ∼1660 cm^-1^ and ∼1690 cm^-1^. The Pearson correlation coefficient (R) was computed and a two-tailed t-test was performed to assess the likelihood of random association between spectral and biochemical data, and considered significant at p<0.05

### 3.5. Correlation between 1660/1690 cm-1 intensity and area ratios and PYD/(DHLNL+HLNL) ratio

Finally, we assessed whether the 1660/1690 cm^-1^ intensity or area ratios were correlated with PYD/(DHLNL+HLNL) content measured by LC-MS. We found positive correlation for both intensity and area ratios (R=0.501, p=0.025 and R=0.571, p=0.009, respectively) suggesting that these spectral ratios are indicative of trivalent over divalent collagen cross-links and hence collagen maturity.

## 4. DISCUSSION

Collagen crosslinks add stability to the organic matrix, preventing micro-fibrils from sliding over each other. Maturation of collagen cross-links is important to ensure resistance to crack growth but also bone strength ^(3,25,26)^. In the present study, we proposed a new method of decomposition of the amide I band by FTIR spectroscopy that takes into account secondary structure of collagen. Although our method to assign subbands was based on second derivative, we employed more subbands than previously reported to account for secondary structure of type I collagen ^(12,15)^. Despite this modification of the original method developed by Paschalis and collaborators, we evidenced that intensities and areas of ∼1660 cm^-1^ and ∼1690 cm^-1^ subbands were still positively correlated with the PYD and DHLNL/HLNL content of the collagen matrix, determined by LC-MS, but also the gene expression level of *Plod2* and *Lox*, two critical enzymes involved in collagen cross-link formation. Furthermore, changes in intensities and areas of the ∼1660 cm^-1^ subband following UV exposure mimicked changes in PYD content determined by LC-MS. As expected, changes in intensities and areas of the ∼1690 cm^-1^ subband after acid treatment also mirrored changes in DHLNL/HLNL content determined by LC-MS. These results are in agreement with previous studies where the ∼1660 cm^-1^ was found correlated to PYD content and similar to changes reported after UV exposure ^(12,13)^. Interestingly, we evidenced a strong correlation between intensity and area of the ∼1680 cm^-1^ subband and DPD content measured by LC-MS in agreement with previous observation of Paschalis et al. ^(13)^. However, we did not observe a reduction in this subband intensity or area after UV exposure as one could expect. Furthermore, intensity and area of the ∼1680 cm^-1^ subband were not correlated with the gene expression level of *Plod2* and *Lox* genes. The ∼1680 cm^-1^ location has been shown to be a signature of advanced glycation end product but also of parallel β-sheet ^(15,17)^. The contribution of DPD to this underlying subband with our method remains to be fully validated as the correlation we observed between spectral and biochemical data might just be due to chance. This also imply that the ∼1660/1680 cm^-1^ ratio is not a good marker of the extent of PYD and DPD in bone tissue.

However, the intensity and area ratios ∼1660/1690 cm^-1^ correlated PYD/(DHLNL+HLNL) significantly with the biochemical ratio, suggesting that our method accurately decomposed the spectral signature of PYD and divalent collagen cross-links from secondary structure. This correlation is also in agreement with the seminal observations made by Paschalis et al and suggest that when curve fitting is performed correctly, FTIR microspectroscopy is a valid method for assessing the extent of enzymatic collagen cross-linking. In opposition to LC-MS, FTIR microspectroscopy is non-destructive and allows an accurate assessment of enzymatic collagen cross-linking and distribution in bone tissue sample. This is of interest as this methodology allows for investigation of cross-links heterogeneity in a tissue that could not be obtained by other technology.

Of note, we did not observe dramatic improvement of any correlation between intensity or area ratio with biochemical data. These observations suggest that both intensity and area ratio could be used for determination of trivalent and divalent collagen cross-links in bone tissue section. This is not so surprising as in transmission mode, FTIR signal follows the Beer’s law that uses intensity rather than area to link absorbance to concentration of an analyte. However, it is important to keep in mind that the ∼1660 cm^-1^ and ∼1690 cm^-1^ subbands are not direct measurement of PYD, DHLNL and HLNL but rather are a reflect of the perturbation these cross-links exert on the carbonyl groups of collagen molecule. This is exemplified by lower intensity of the ∼1690 cm^-1^ intensity as compared with the ∼1660 cm^-1^ subband despite higher content of divalent cross-links than PYD measured by LC-MS. The originality of our study also consists in a double validation against LC-MS gold standard but also gene expression of key enzyme involved in collagen cross-linking. This approach of correlating gene expression levels with spectral data has not yet been used for bone tissue, but was employed for breast tissue ^(27)^.

In summary, we developed and validated in the present study a new method of amide I decomposition that takes into account secondary structure of collagen molecule and yet allows for accurate determination of PYD and enzymatic divalent cross-links of type I collagen in bone tissue section. This method confirms previous report that the ∼1660 cm^-1^ and ∼1690 cm^-1^ subbands are indicative of PYD and divalent cross-link content. This method of spectral processing allows for investigation of tissue content and distribution of collagen cross-linking.

## 5. AUTHOR CONTRIBUTION

**Aleksandra Mieczkowska:** Investigation, Formal analysis and **Guillaume Mabilleau:** Conceptualization, Investigation, Formal analysis, Writing -Review & Editing, Supervision, Funding acquisition, Data curation

## 6. ACKNOWLEDGEMENTS

We are thankful to Prof Peter Gardner (University of Manchester) for supplying the Mie scattering correction routine for Matlab. We are also grateful to the institutional animal lab SCAHU, SFR ICAT 4208, Univ Angers, for their help with animal care. This project was funded by an institutional grant from the University of Angers.

## 8. DATA AVAILABILITY STATEMENT

The data that support the findings of this study are available from the corresponding author upon reasonable request.

